# The immunophilin Zonda controls regulated exocytosis in endocrine and exocrine tissues

**DOI:** 10.1101/2020.05.28.122465

**Authors:** Rocío de la Riva Carrasco, Sebastián Perez Pandolfo, Sofía Suarez Freire, Nuria M. Romero, Zambarlal Bhujabal, Terje Johansen, Pablo Wappner, Mariana Melani

## Abstract

Exocytosis is a fundamental process in physiology, communication between cells, organs and even organisms. Hormones, neuropeptides and antibodies, among other cargoes are packed in exocytic vesicles that need to reach and fuse with the plasma membrane to release their content to the extracellular milieu. Hundreds of proteins participate in this process and several others in its regulation. We report here a novel component of the exocytic machinery, the *Drosophila* transmembrane immunophilin Zonda (Zda), previously found to participate in autophagy. Zda is highly expressed in secretory tissues, and regulates exocytosis in at least three of them: the ring gland, insulin-producing cells and the salivary gland. Using the salivary gland as a model system, we found that Zda is required at final steps of the exocytic process for fusion of secretory granules to the plasma membrane. In a genetic screen we identified the small GTPase RalA as a crucial regulator of secretory granule exocytosis that operates upstream of Zda in the process.

## INTRODUCTION

Exocytosis is a fundamental cellular process required for delivery of proteins, lipids and carbohydrates to the extracellular milieu, so hormones, antibodies and neuropeptides, among others, are released from the cells where they are produced by this mechanism. The exocytic process requires the genesis of secretory vesicles in which export products are packed. These vesicles sprout from the trans-Golgi network in an immature exocytosis-incompetent state, and thereafter, vesicles undergo maturation, in a process that includes homotypic fusion, condensation and acidification of their content. During this process, incorporation of specific vesicle membrane proteins occurs, including SNARES and Synaptotagmins required for membrane fusion. Then, mature exocytic vesicles or secretory granules (SGs) are directionally transported to the cellular apical domain, where prior to secretion, a series of events that include tethering, priming, triggering and fusion to the plasma membrane take place, each of them executed by specific molecular complexes and their regulators (Sugita, 2007).

Most cellular models for studying exocytosis rely on the analysis of a sole readout: Intracellular accumulation of SGs and/or exocytosis of SG content. The salivary gland of *Drosophila melanogaster* larvae is a useful model to study the mechanisms involved in exocytosis (Biyasheva, Do, Lu, Vaskova, & Andres, 2001; Tran & Ten Hagen, 2017). At late 3^rd^ larval instar, salivary glands synthesize a series of mucins called Glue proteins that are packed into SGs, known as Glue granules (GGs). Immature GGs initially sprout from the trans-Golgi network (TGN) as 1μm diameter vesicles, reaching then a mature size of around 5μm after several events of homotypic fusion and maturation (Jason Burgess et al., 2011; Reynolds, Zhang, Tran, & Ten Hagen, 2019). At the onset of puparation GGs undergo exocytosis and Glue proteins are released to the salivary gland lumen, from where they are later secreted to adhere the puparium to the substratum (Biyasheva et al., 2001; Costantino et al., 2008). Recently, the final steps of GG exocytosis at the salivary gland were described in detail. Fusion of the GG to the Apical Plasma Membrane (APM) results in transfer of the lipid phosphatidylinositol 4,5-bisphosphate (PI(4,5)P_2_) from the APM to the GG membrane, an event that enables recruitment of the small GTPase Rho1 to the GG, followed by simultaneous activation of Diaphanous (Dia) and Rho-associated kinase (Rok) on the GG membrane. These events result in polymerization of an acto-myosin mesh around the GGs, which is critical for efficient release of GG content into the salivary gland lumen (Rousso, Schejter, & Shilo, 2016; Tran, Masedunskas, Weigert, & Ten Hagen, 2015). However, our knowledge on how GGs fuse with the plasma membrane remains incomplete.

FK506-binding proteins (FKBPs) are immunophilins that can bind the immunosuppressant KF506, and display peptidyl-prolyl *cis/trans* isomerase activity (PPIase). FKBPs participate in a myriad of cellular activities, including protein folding, receptor signaling and transcription (Ghartey-Kwansah et al., 2018; Kang, Hong, Dhe-Paganon, & Yoon, 2008). FKBP8 is a non-canonical member of this family, as its PPIase activity depends on binding to Ca^++^-conjugated Calmodulin, and includes a transmembrane domain on its carboxy-terminus, a unique feature among FKBPs (Bhujabal et al., 2017; Kang et al., 2008). FKBP8 has been reported to interact with several proteins such as Bcl-2, Bcl-XL (Y. Chen, Sternberg, & Cai, 2008; Haupt et al., 2012), HSP90 (Okamoto et al., 2006; Shimamoto et al., 2014), Rheb (Bai et al., 2001), PDH2 (Barth et al., 2007) and LC3 (Bhujabal et al., 2017). In this way, FKBP8 regulates diverse cellular processes, including apoptosis, mitophagy and hypoxic responses. Previously, we have characterized the function of Zonda (Zda), the predicted *Drosophila* FKBP8 ortholog, as a key regulator of early steps of autophagy (Melani et al., 2017).

In this work we report a novel function for the transmembrane immunophilin Zda in regulated exocytosis in different secretory tissues, including the prothoracic gland (PG), insulin producing cells (IPCs) of the brain, and salivary glands. Detailed analysis of Zda loss-of-function phenotypes at the salivary gland revealed that it is required for GG fusion with the plasma membrane, but not for their biogenesis or maturation. Through a genetic screen of components known to participate in secretory granule fusion to the plasma membrane, we identified the small GTPase RalA as a key player in GG exocytosis, which genetically interacts with Zda.

## RESULTS

### Zonda is expressed in secretory tissues

We made use of the allele *zda*^*trojan*^ to gain insights into the tissues in which *zda* is expressed. *zda*^*trojan*^ insertion generates a truncated product (Supplementary Figure 1A), and therefore renders a null allele that is lethal in homozygosis, which does not complement with another previously characterized *zda* null allele (*zda^null^)* (Melani et al., 2017). Moreover, expression of a UAS-*mCh-zda* construct driven by *zda*^*trojan*^ completely restored viability, strengthening the notion that *zda*^*trojan*^ is a genuine loss-of-function allele, and that mCh-Zda is fully functional. Interestingly, overexpression of truncated versions of mCh-Zda, lacking its transmembrane domain (mCh-Zda^Δ™^) or its calmodulin/TPRdomain (mCh-Zda^ΔCaM/ΔTPR^), failed to rescue lethality of the *zda*^*trojan*^ allele, indicating that these domains are required for Zda function (Supplementary Figure 1B,C).

*zda*^*trojan*^ contains a gene trap cassette, derived from a MiMIC insertion in *zda’s* second intron (Supplementary Figure 1A). The cassette includes a Trojan GAL4 exon composed of a splice acceptor, a T2A peptide, a GAL4 coding sequence and an Hsp70 transcription termination signal (Diao et al., 2015). We crossed this allele with a UAS-mCD8-GFP reporter, and observed that *zda* is highly expressed in glandular tissues and secretory cells, among other organs, as previously reported by high-throughput anatomy RNA-seq data (Gelbart, W.M., Emmert, 2013). We detected strong expression in the salivary glands, the ring gland (RG), the lymph gland, insulin producing cells (IPC) of the brain, a subset of cells of the intestine, and secondary cells of the adult male accessory gland (Figure 1A-F). The fat body also displayed high expression levels, as also did differentiated cells of the eye imaginal disc, posterior to the morphogenetic furrow (Figure 1 G, H). Other imaginal discs, such as the wing disc showed no significant expression of *zda*^*trojan*^ under the same imaging conditions (Figure 1I). The ejaculatory duct also displayed high expression levels (Figure 1J), unlike the testis where expression was minimal (Figure 1K), while no significant expression was observed in adult ovaries (Figure 1L). Since *zda* is expressed at high levels in several tissues or cell types with secretory function, we sought to explore a possible role of Zda in exocytosis.

**Figure 1.**
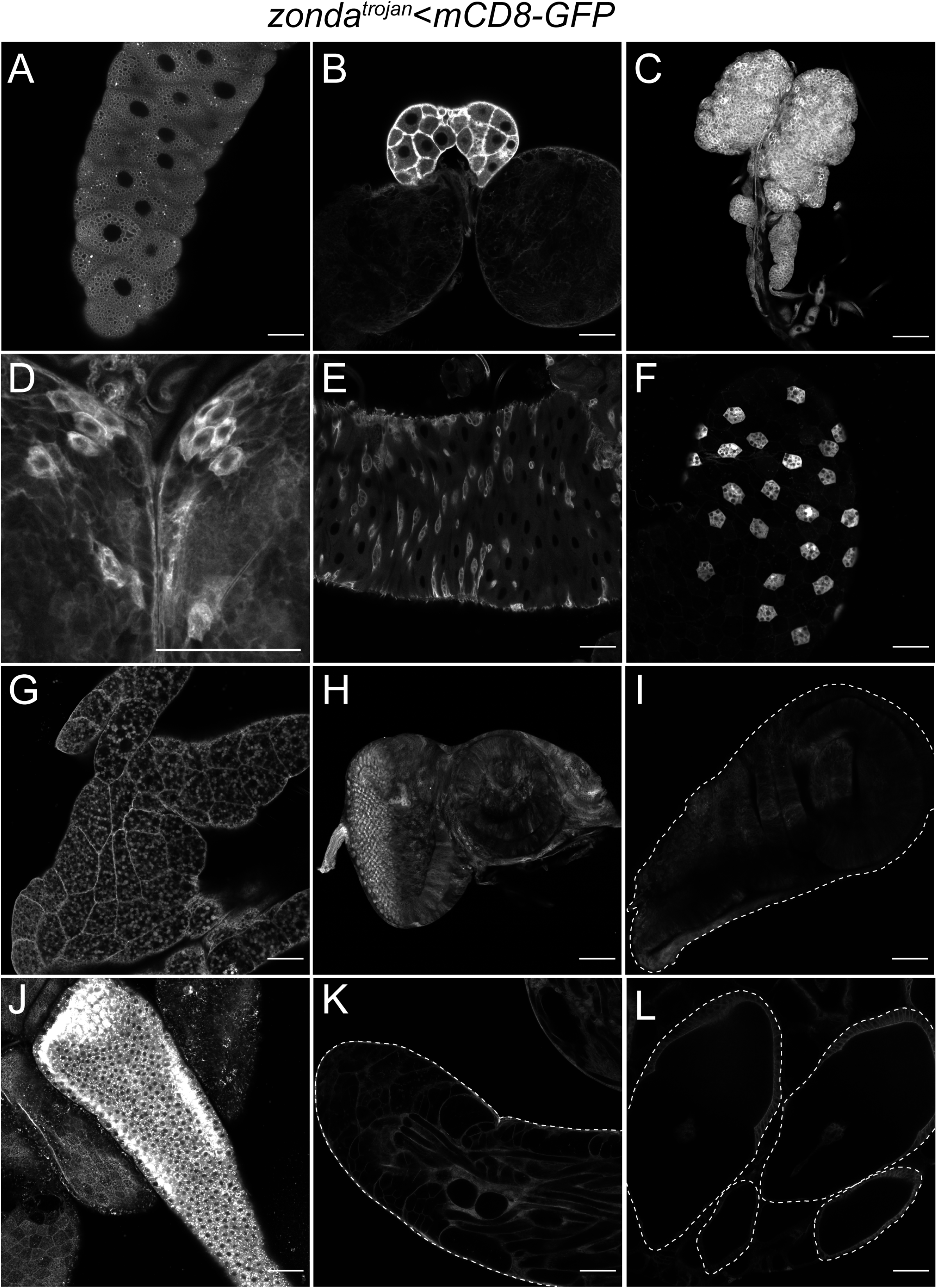
Zonda is expressed in secretory tissues. The *zda*^*trojan*^ line was crossed to UAS-mCD8-GFP flies, and tissues were dissected and observed directly under the confocal microscope. Larval salivary gland (A), ring gland (B), brain (C), lymph gland (D), intestine (E), fat body (F), eye imaginal disc (G), wing imaginal disc (H), adult male accessory gland (I), ejaculatory duct (J), testis (K), adult female egg chambers (L). For comparative purposes all images were acquired using the same microscope set up.

### Zonda is required for exocytosis of ecdysone, Insulin-like peptide 2 and Glue proteins

We hypothesized that the *zda* expression pattern might reflect a function of Zda in secretion. We therefore analyzed Zda function in the RG, IPCs and salivary glands. The RG is composed of three different cell types specialized in secretion of specific products. The prothoracic gland (PG) encompasses most of the RG, and is dedicated to biosynthesis and secretion of the steroid hormone 20-hydroxy ecdysone (20E) (Yamanaka, Marqués, & O’Connor, 2015). Also, PG cells can be readily distinguished from other RG cell types as they are innervated by axons of PTTH-producing neurons. As depicted in Figure 2A, *zda* is highly expressed in PG cells that are innervated by PTTH axons. To test if Zda might be involved in 20E exocytosis, we expressed a *zda* RNAi using the PG driver *phantom*-Gal4, and observed a significant delay in pupariation time as compared to controls (Figure 2B). Supplementation of the culture medium with 20E resulted in complete rescue of normal pupariation time (Figure 2B), suggesting that Zda knock down in PG cells results in reduced levels of circulating 20E (Yamanaka et al., 2015).

**Figure 2.**
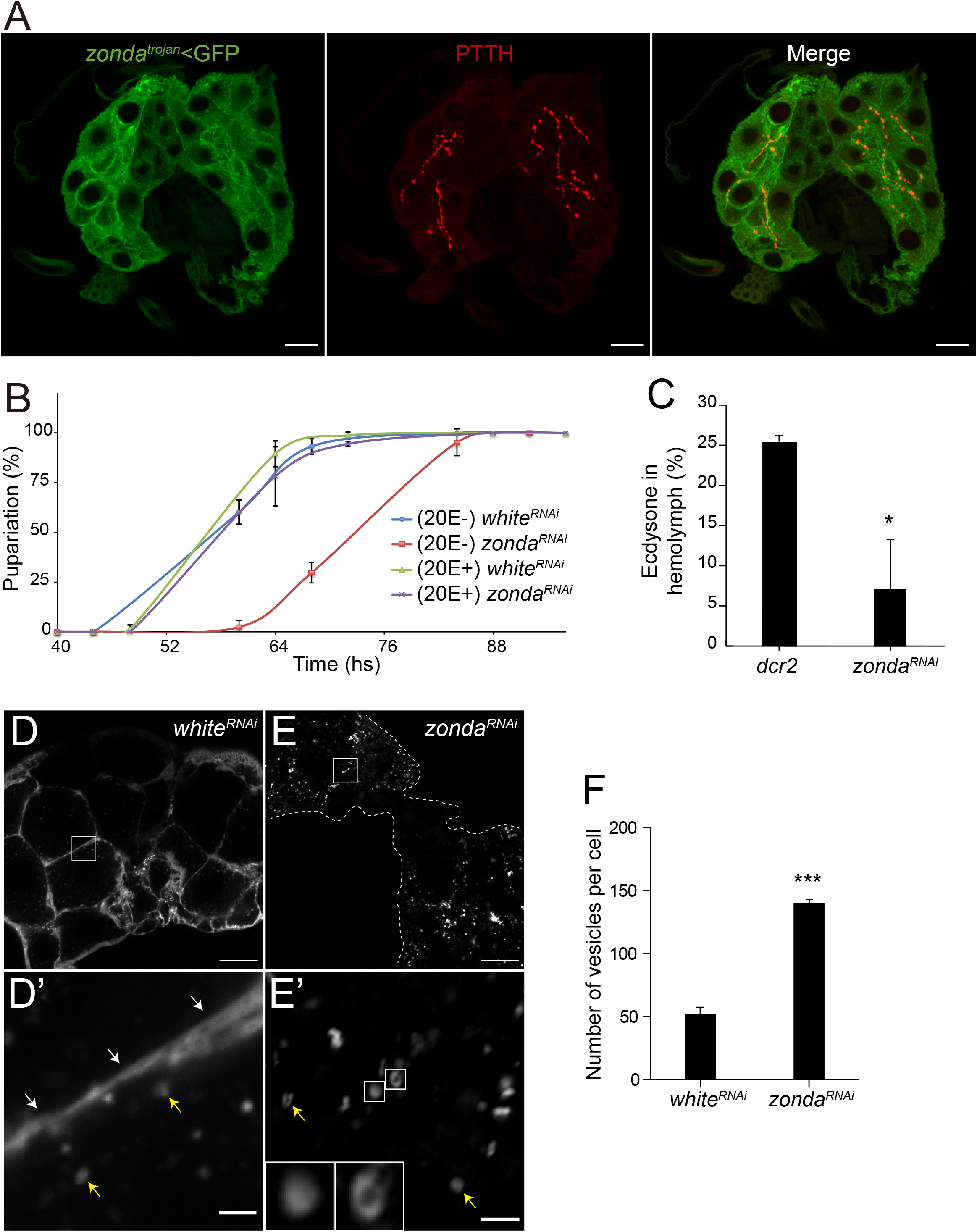
Zonda is required for ecdysone exocytosis at the prothoracic gland. (A) *zda*^*trojan*^ is expressed at high levels in the prothoracic gland (PG), as revealed by visualization of mCD8-GFP (green) that coincides with PPTH-labeled axons (red). (B) Puparation time was recorded in control (*phantom*<*white*^*RNAi*^) and Zda knock-down larvae (*phantom*<*zda^RNAi^*) that were grown either in control media or media supplemented with ecdysone (20E). (C) Quantification by ELISA of 20E levels in hemolymph relative to levels of total 20E in L2 larvae homogenates; * p<0.05. (D-E) Confocal images of wandering larvae ring glands that express Synaptotagmin-1-GFP (green) under control of *P0206*-*Gal4*. (D) Control ring glands (UAS-*white*^*RNAi*^) and (E) Zda knock-down ring glands (UAS-*zda^RNAi^*) are compared. Insets: (D’) Synaptotagmin is mostly concentrated at the plasma membrane in control individuals (white arrows), while the number of intracellular vesicles labeled with syt-1-GFP is low (yellow arrows); (E’) in Zda knock-down larvae many syt-1-GFP vesicles can be observed inside PG cells (yellow arrows); the insets below show magnified images of the vesicles boxed in panel E’. Scale bars: D, E = 20μm, D’, E’ = 2μm. (F) quantification of Syt-1 positive vesicles detected in each genotype. N = 3 for each genotype. ***p<0,001.

To investigate if the above Zda loss-of-function phenotype may arise from impaired 20E secretion, we determined the levels of circulating 20E relative to the total 20E larval content, and found that Zda silencing in the PG results in 3.5 times reduction of circulating 20E (Figure 2C). It was recently shown that after being synthesized in PG cells, ecdysone is packaged in secretory vesicles, and released to the extracellular milieu by exocytosis (Yamanaka et al., 2015). As these vesicles are Synaptogamin-1 (Syt-1) positive, we analyzed the presence of Syt-1-GFP vesicles in PG cells of control *versus zda* RNAi-expressing wandering larvae. In control individuals, we observed few intracellular Syt1-GFP positive vesicles as previously reported (Yamanaka et al., 2015), with most of the Syt-1-GFP label localized at the plasma membrane (Figure 2D, F), suggesting that most vesicles have already exocytosed their content. In contrast, in Zda-deficient PG cells, Syt-1-GFP-positive vesicles could be readily detected, and concomitantly almost no Syt-1-GFP was observed at the plasma membrane (Figure 2E, F), suggesting that exocytosis is impaired in these PGs. We thus conclude that Zda is required at the PG for exocytosis of Syt-1-positive ecdysone-containing vesicles.

To further evaluate a possible role of Zda in regulated exocytosis, we turned to the IPCs. IPCs are brain neurosecretory cells clustered in two groups of 7 cells each that produce and secrete insulin-like peptides (dILPs), which are released to the hemolymph in response to hormonal or environmental stimuli, and regulate body growth (Cao et al., 2014). We first confirmed that *zda* is highly expressed in IPCs, as its expression colocalizes with that of Dilp2 (Géminard, Rulifson, & Léopold, 2009)(Figure 3A). Next, we tested if Zda is required for dILP secretion by knocking down its expression in IPCs with a *dilp2*-Gal4 driver. This silencing resulted in significant reduction of pupal volume (Figure 3B), which is characteristic of diminished levels of circulating dILPs. Consistent with this, dILP2 levels were significantly higher inside the IPCs in well-fed *zda*^*RNAi*^ larvae, as compared to wild type controls in which dILP2 was detected at high levels in IPCs of starved, but not of well-fed larvae (Figure 3C, D). These results suggest that Zda is required at the IPCs for dILP2 exocytosis.

**Figure 3.**
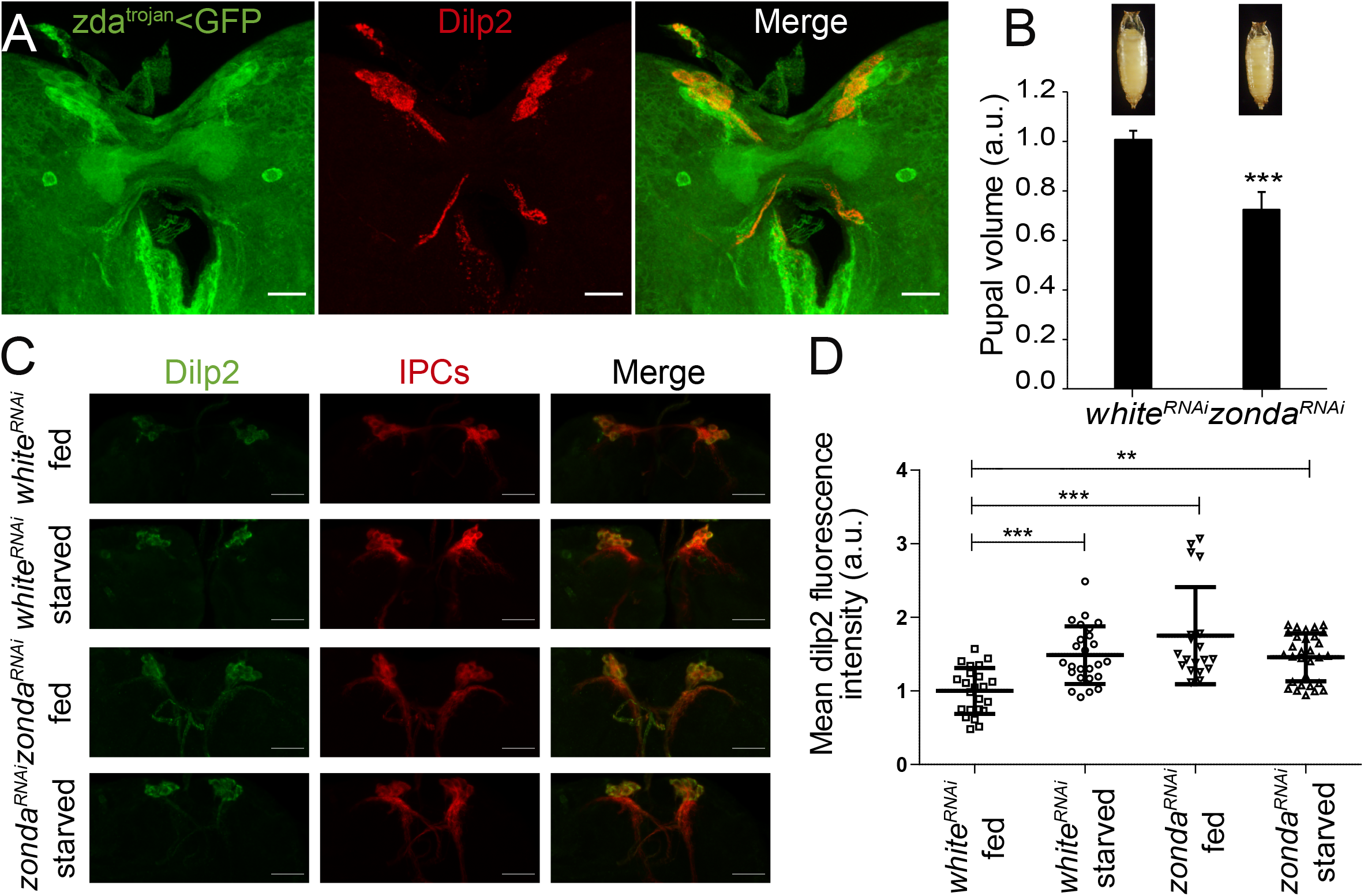
Zonda is required in Insulin Producing Cells for Dilp2 exocytosis. (A) *zda*^*trojan*^ expression is high in IPCs, as revealed by mCD8-GFP expression (green), and colocalization with Dilp2 (red). (B) Downregulation of *Zda* in IPCs (dilp2<*zda*^*RNAi*^) provokes reduction of pupal volume compared to control pupae (dilp2<*white*^*RNAi*^). Representative images are shown. Control: N = 85; *zda*^*RNAi*^: N = 65. (C, D) Zda downregulation provokes accumulation of Dilp2 in IPCs even under feeding conditions; the Dilp2 signal in IPCs of 3^rd^ instar larvae upon 16 hours starvation was compared to that of fed individuals. (C) Dilp2 was detected by immunofluorescence (green), while IPCs where identified by expression of UAS-cherry under the control of Dilp2-Gal4 (red). Representative images are shown. (D) Quantification of the average fluorescence intensity in the experiment of panel (C). *white*^*RNAi*^ fed N = 24; *white*^*RNAi*^ starved N = 24; *zda*^*RNAi*^ fed N = 19; *zda*^*RNAi*^ starved N = 30. ** p<0,01; *** p<0,001. Scale bar = 50 μm.

Finally, we looked at exocytosis of Glue granules (GGs) in larval salivary glands. GGs are exocytic vesicles packed with mucins named Salivary Gland Secretion (SGS) that serve to adhere the animal to the substratum at the time of puparation (Biyasheva et al., 2001). GGs form in response to an ecdysone peak at the onset of the larval wandering stage, and are exocytosed in response to a later ecdysone peak at the time of puparium formation (Biyasheva et al., 2001). To evaluate exocytosis of GGs, we utilized a transgenic line that expresses one of the SGSs, SGS3, fused to GFP (SGS3-GFP) (Biyasheva et al., 2001). In wandering larvae, salivary glands normally contain large amounts of SGS3-GFP (Figure 4A, A’), while after pupariation, SGS3-GFP is exocytosed to the lumen of the gland and released outside of the puparium (Figure 4B, B’, E). When Zda was downregulated in salivary glands, we found normal levels of SGS3-GFP in salivary glands of wandering larvae, indicating that the mucin is normally produced (figure 4C, C’), while at the prepupal stage SGS3-GFP failed to be secreted, and remained inside the gland (Figure 4D, D’, E). These observations suggest that Zda is required at the salivary gland for exocytosis of GGs. To analyze if this is the case, we dissected salivary glands of control and Zda knock down prepupae, and observed that, whereas control salivary glands do not contain GGs (Figure 4F), in Zda knock down salivary gland cells, a large number of GGs were present (Figure 4G). These results indicate that Zda is required for GG exocytosis in salivary glands, and more generally, that Zda regulates exocytosis in exocrine and endocrine glands.

**Figure 4.**
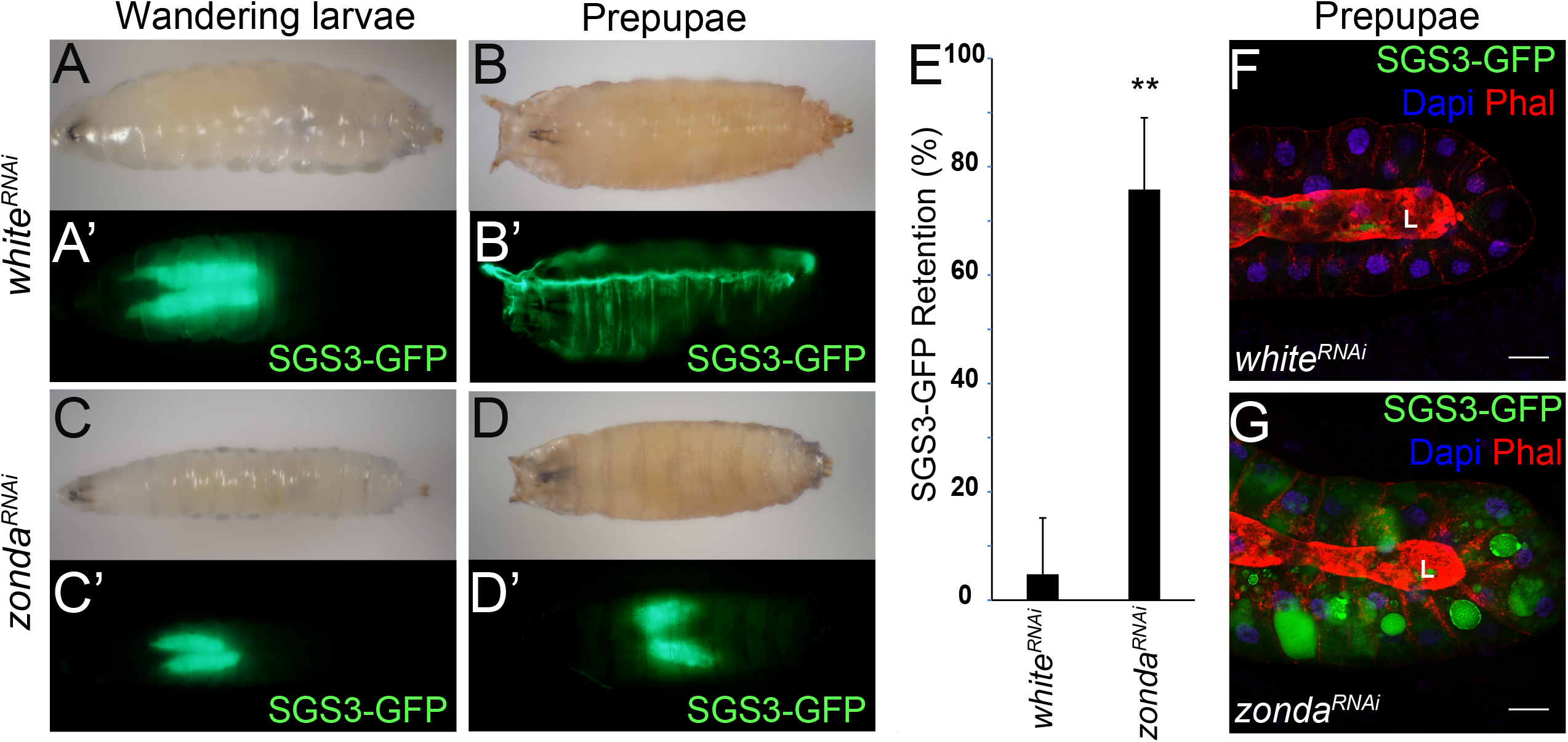
Zonda is required in salivary glands for Glue granule exocytosis. (A-D) Secretion of Sgs3-GFP (green) in control and Zda-knock down larvae and prepupae. Larvae of both genotypes accumulate comparable levels of Sgs3-GFP in their salivary glands (A, A’ and C, C’). After puparation, control larvae have secreted Sgs3-GFP, which is extruded outside the puparium (B, B’), while *zda*^*RNAi*^ prepupae retain Sgs3-GFP inside their salivary glands (D, D’). (E) quantification of Sgs3-GFP retention inside salivary glands in prepupae; N = 192 for *white*^*RNAi*^ and N = 133 for *zda*^*RNAi*^. ** p<0,01. (F, G) Sgs3-GFP is retained inside salivary gland cells in Zda-knock down prepupae. Confocal images of control (*white*^*RNAi*^) and Zda-knock down (*zda*^*RNAi*^) prepupal salivary glands; Sgs3-GFP-labeled Glue granules (green), nuclei stained with DAPI (blue), and phalloidin (red). “L” indicates the lumen. Scale bar: 50μm

### Zonda is required for GG fusion with the plasma membrane

We utilized the salivary gland to study in more detail the role that Zda plays in exocytosis. GGs emanate from the Trans Golgi Network as 1 μm vesicles, and reach a mature size of 5 μm prior to fusion with the APM (Jason Burgess et al., 2011). We compared GG diameter in salivary glands of Zda knock down and control wandering larvae, just prior to the stage when exocytosis is expected, and observed that GG size was not altered (Figure 5A-C), suggesting that Zda is not required for GG biogenesis or maturation.

**Figure 5.**
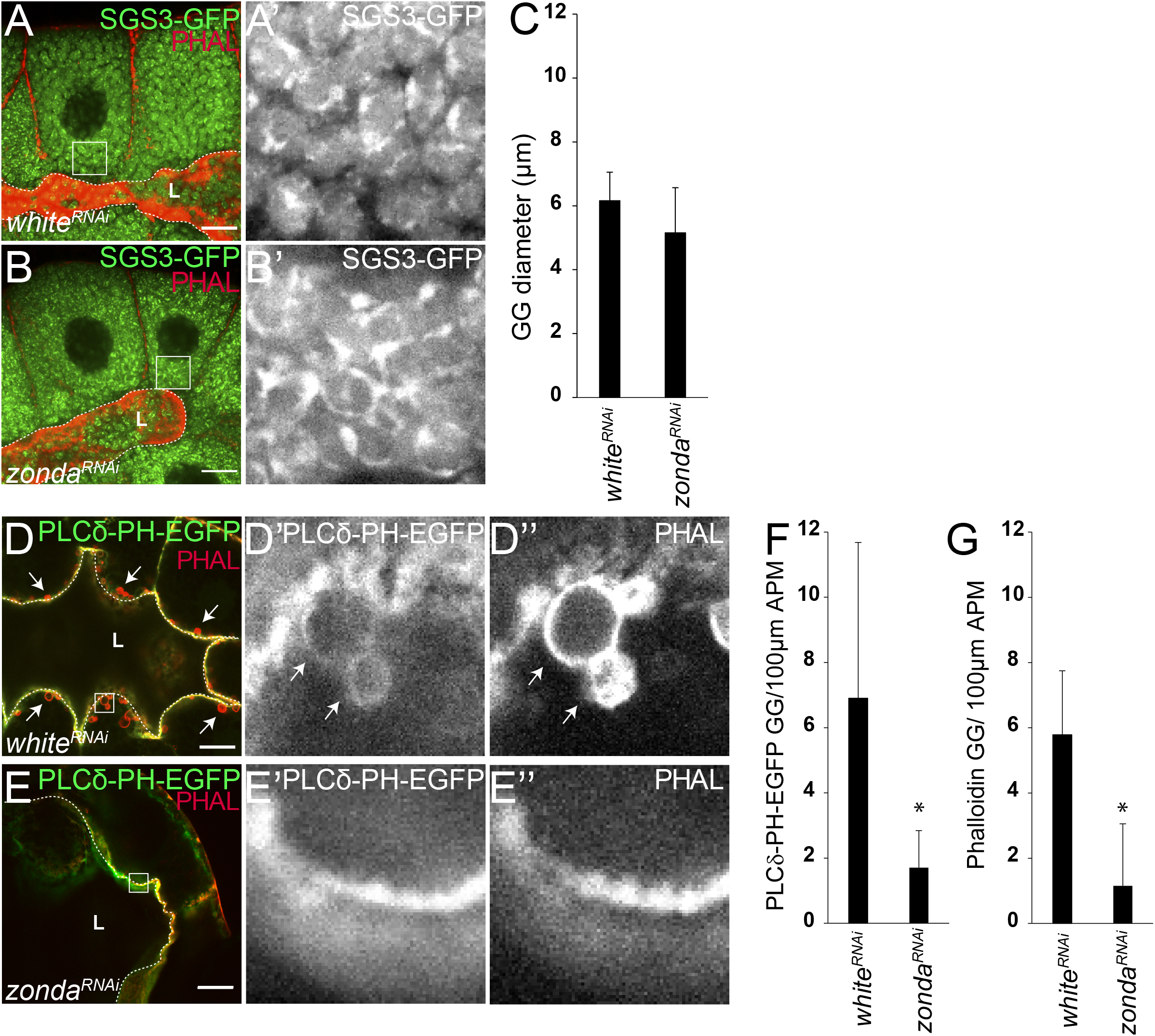
Zonda is required for Glue granule (GG) fusion with the plasma membrane. (A-C) Zda is not required for GG biogenesis or maturation. (A, B) Confocal images of salivary glands dissected from wandering larvae of control (*white*^*RNAi*^) (A, A’) and *zda*^*RNAi*^ (B, B’) individuals prior to puparation expressing Sgs3-GFP (green), and labelled with phalloidin (red). “L” indicates the lumen that is also marked with a dashed line. Scale bar: 20μm. Crop size 20μm×20μm. (C) Quantification of GG diameter in both genotypes; no statistical difference was detected; 40 GGs were scored for each genotype (N = 4). (D-G) Zda is required for GG fusion with the PM. (D-E) Confocal images of wandering larvae salivary glands expressing PLCδPH-EGFP (green) and stained with phalloidin (red). While in control larvae GGs are positive for PLCδPH-EGFP (D, D’), and stain positive for phalloidin (D, D”), *zda*^*RNAi*^ GGs are negative for both markers (E-E”). Arrows mark GGs positive for PLCδPH-EGFP and phalloidin. “L” indicates the lumen that is also marked with a dashed line. Scale bar: 20μm. Crop size: 15μmx15μm. (F, G) Quantification of GGs containing PLCδPH-EGFP (F) or phalloidin (G) in 100 μm of plasma membrane. N= 40 for each genotype. Statistically significant differences were found for the two markers analyzed (* p<0,05).

PI(4,5)P_2_ is a lipid of the inner leaflet of the plasma membrane, absent from intracellular organelles (Phan et al., 2019). Upon fusion of mature GGs with the APM, PI(4,5)P_2_ incorporates to the membrane of the GG, and this lipid transfer can be followed by looking at the PI(4,5)P_2_ reporter PLCδPH-EGFP (Rousso et al., 2016; Tran et al., 2015). GG fusion to the APM triggers the formation of a filamentous actin mesh around the GG that can be readily detected by life-act-Ruby or phalloidin (Rousso et al., 2016; Tran et al., 2015), so that only mature GGs that successfully fuse with the APM are positive for PI(4,5)P_2_ and actin markers (Figure 5D-F, G). Both PI(4,5)P_2_ and actin recruitment to GGs was significantly reduced in Zda knock down larvae (Figure 5E-G), suggesting that Zda is required for fusion of GGs to the APM.

### RalA is required for GG fusion with the plasma membrane upstream of Zonda

Given that Zda is not required for GG biogenesis or maturation, but necessary for fusion of GGs to the plasma membrane, we hypothesized that Zda may participate in docking, priming, triggering or fusion of GGs to the APM, so we performed a loss-of-function screen aimed at identifying genes that participate in these processes, which may cooperate with Zda in exocytosis of GGs. We focused particularly on genes highly expressed at the salivary gland, according to high throughput data compiled at flybase (flybase.org). We expressed in salivary glands double stranded RNAs or dominant negative alleles of candidate genes, which included Rabs (Pfeffer, 2017), SNAREs (Zorec, 2018), the subunits of the exocyst complex (Heider & Munson, 2012), the small GTPase RalA (Shirakawa & Horiuchi, 2015), Synaptotagmins (Gustavsson & Han, 2009), AP proteins (Azarnia Tehran, López-Hernández, & Maritzen, 2019), RE-PM contact site proteins (Saheki & De Camilli, 2017), and calcium channels (Rizo & Rosenmund, 2008) (Supplementary Table 1). As a read out for the screen, we looked at SGS3-GFP retention in prepupae (Figure 4). Loss of function of 18 out of the 64 genes analyzed provoked retention of SGS3-GFP with a penetrance of 50% or higher (Supplementary Table 1 and Supplementary Figure 2).

The retention phenotype after suppressing the activity of these 18 candidate genes was further analyzed by confocal microscopy, specifically by looking at GG size and actin polymerization around the GGs. Out of the 18 candidates, loss of function of 15 of the genes resulted in blockage of GG biogenesis and/or maturation, since no GGs or very small GGs were detected (Supplementary Figure 3A, B). Among these 15 genes were the eight subunits of the exocyst complex; the small GTPase Rab1, known to mediate dynamic membrane trafficking between ER and Golgi (Plutner et al., 1991); the adaptor proteins Ap-1-2β and the Arf GEF Sec71, previously reported as essential for GG biogenesis (Jason Burgess et al., 2011; Torres, Rosa-Ferreira, & Munro, 2014); and the syntaxins Syx5 and Syx7, Sec20, and Syt4 (Supplementary Figure 3E). On the other hand, two of the hits, Rab11 and EpsinR, resulted in mature GGs that appeared covered with filamentous actin, although mislocalized at the basolateral domain of the cell (Supplementary Figure 3C, E). Finally, one remarkable hit was the small GTPase RalA, whose loss of function resulted in GGs of mature size (Figure 6A-C; Supplementary Figure 3D, E) that never fuse with the plasma membrane, do not contain PI(4,5)P_2_, and are not surrounded by polymerized actin (Figure 6D-G). Given the similarities between Zda and RalA knock down phenotypes, we analyzed possible genetic interactions between the two genes. Over-expression of Zda led to partial rescue of the RalA knock-down phenotype (Figure 6H), suggesting that Zda functions downstream of RalA in GG exocytosis.

**Figure 6.**
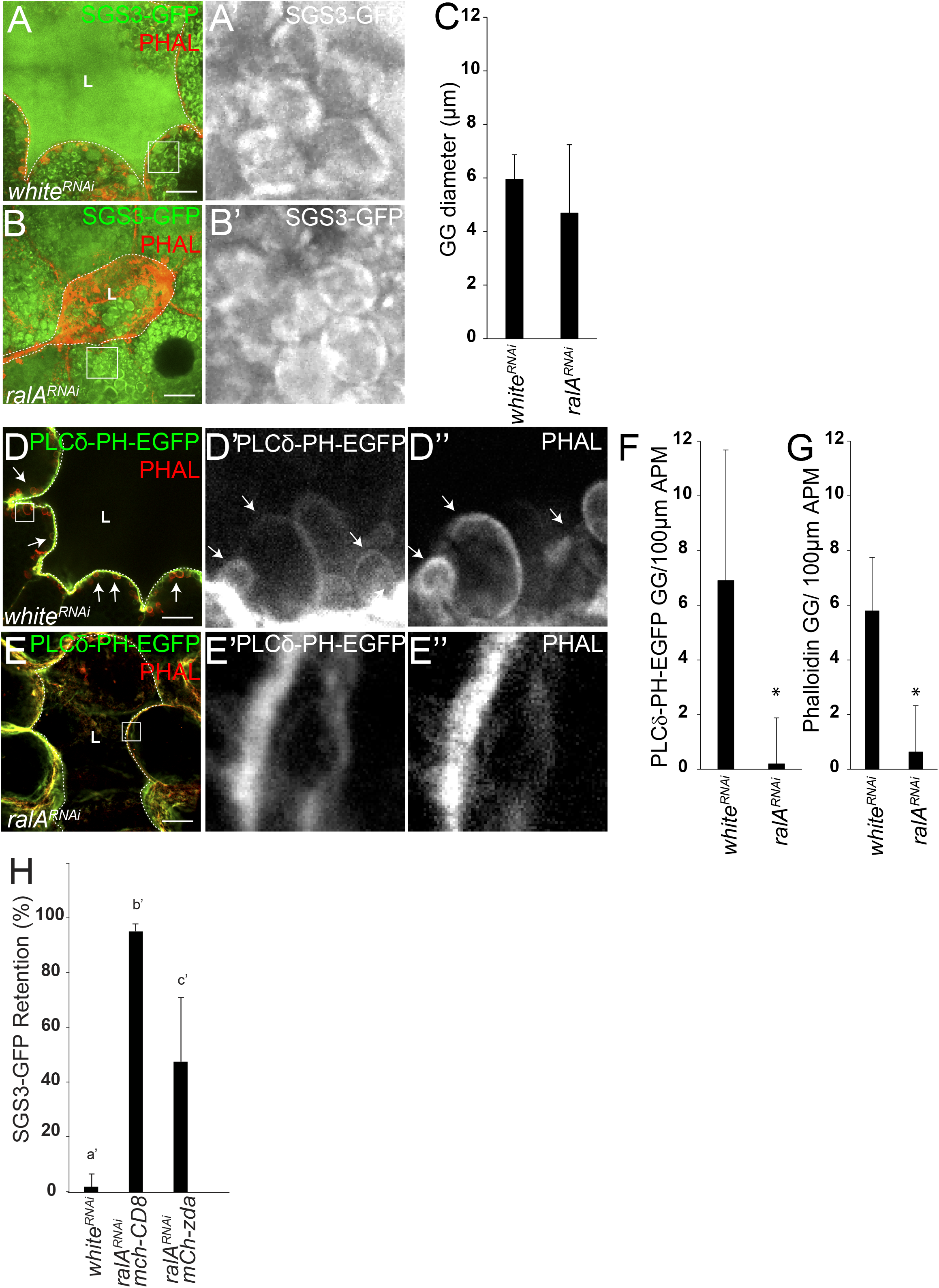
RalA is required for Glue granule fusion with the plasma membrane. (A-C) RalA is not required for GG biogenesis or maturation. (A-B) Confocal images of salivary glands dissected from wandering larvae of control (*white*^*RNAi*^) (A) and *ralA^RNAi^* (B) genotypes expressing Sgs3-GFP (green), and stained with phalloidin (red). “L” indicates the lumen that is also marked with a dashed line. Scale bar: 20μm. Crop size 20μm×20μm. (C) Quantification of GG diameter in both genotypes; no statistical difference was detected; 40 GGs were scored per genotype (N = 4). (D-G) RalA is required for GG fusion with the PM. (D-E) Confocal images of wandering larvae salivary glands expressing PLCδPH-EGFP (green) and stained with phalloidin (red). While in control larvae GGs are positive for PLCδPH-EGFP (D, D’) and phalloidin (D, D”), in *ralA^RNAi^* individuals GGs are negative for both markers (E-E”). Arrows in D-D”point at GGs positive for PLCδPH-EGFP and phalloidin. “L” indicates the lumen that is also marked with a dashed line. Scale bar: 20μm. Crops size: 15μmx15μm. Quantification of GGs containing PLCδPH-EGFP (F) or phalloidin (G) in 100 μm of plasma membrane; N = 40 for each genotype. Statistically significant differences were found for the two markers analyzed (* p<0.05). (H) Genetic interaction between Zda and RalA: The retention phenotype of Sgs3-GFP in salivary glands was scored in control prepupae (*white*^*RNAi*^, N = 196), *RalA^Rnai^* (N = 125) and *RalA^Rnai^* individuals that overexpressed mChZda (N = 102). The RalA loss-of-function phenotype was largely suppressed by overexpression of mChZda. ANOVA test followed by Tukey test was performed; different letters indicate statistically significant differences (p<0,05).

Overall, we have identified Zda as an important player in the process of regulated exocytosis, executing its action at the final steps of the process, just prior to fusion of secretory granules with the APM, likely cooperating in this process with the small GTPase RalA (Figure 7).

**Figure 7.**
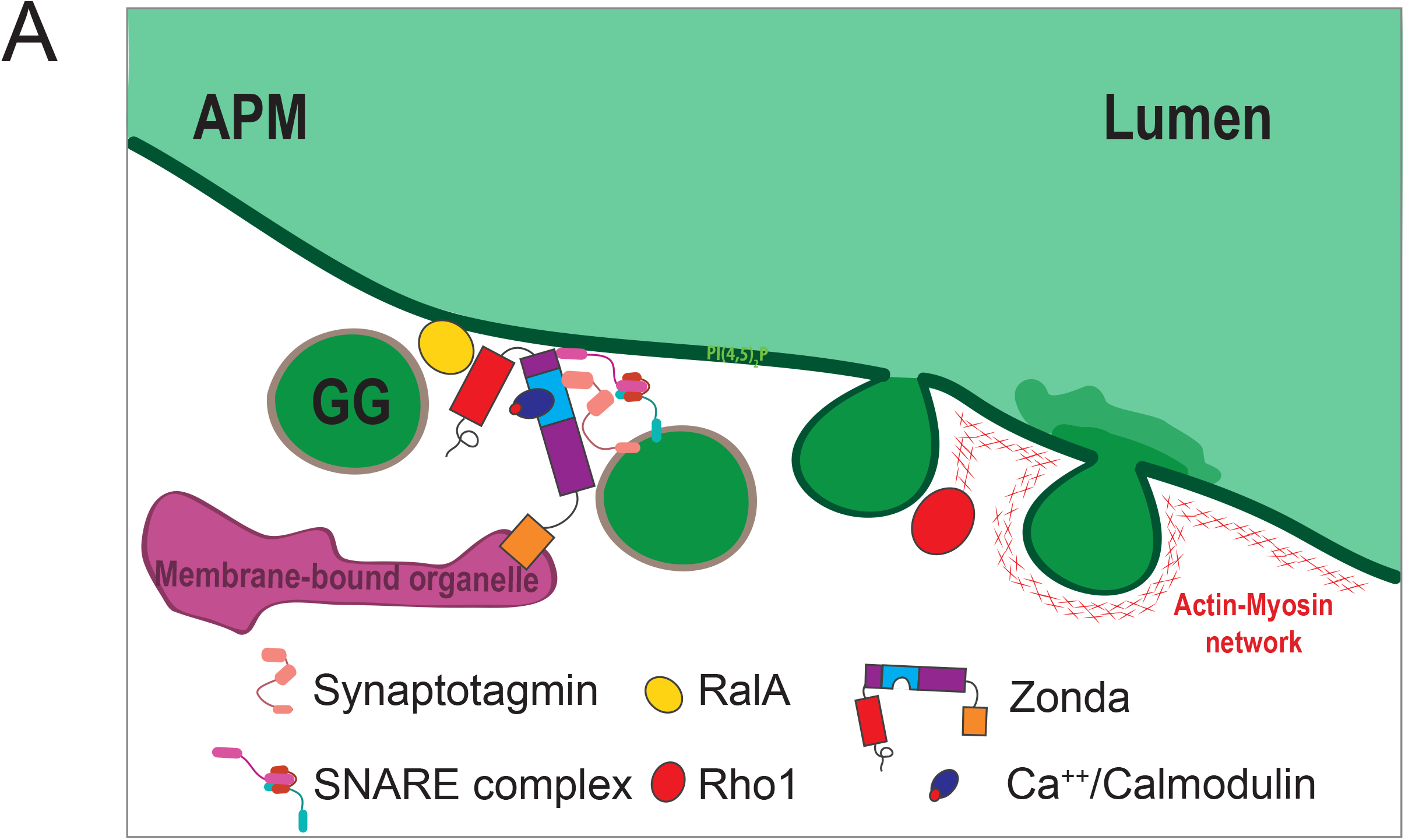
Proposed model of Zonda action in Glue granule exocytosis. Zda operates at final stages of GG exocytosis downstream of RalA, and before GG-APM fusion. Ca^++^/CaM-bound Zda might act as a platform for fusogenic factors, such as SNAREs and Synaptotagmin. Rho-1-induced acto-myosin recruitment to GGs occurs down-stream of Zda action.

## DISCUSSION

In this work we have shown that the *Drosophila* transmembrane immunophilin Zda is highly expressed in secretory tissues, and critically required for regulated exocytosis. In the PG Zda regulates exocytosis of the molting hormone 20E; in IPCs is required for Dilp2 exocytosis, and in salivary glands it controls exocytosis of mucin–containing glue granules. Unlike other genes previously reported to contribute to *Drosophila* salivary gland secretion, such as AP-1 (Jason Burgess et al., 2011), PI4K (J. Burgess et al., 2012), Arl1 and Sec71 (Torres et al., 2014), Hobbit (Neuman & Bashirullah, 2018) and Tango1 (Reynolds et al., 2019), Zda is not required for GG biogenesis, but rather at the final steps of the exocytic process.

Zda, as an immunophilin, encompasses the classical PPI and TRP domains, but in addition includes a Ca^++^/CaM binding domain and a transmembrane domain on its carboxy-terminus (Melani et al., 2017), which we found in this work, are essential for Zda function and *Drosophila* viability. FKBPs are believed to operate as molecular platforms assisting the interaction of components of multi-molecular complexes (Bai et al., 2001; Y. Chen et al., 2008; Haupt et al., 2012; Okamoto et al., 2006; Shimamoto et al., 2014). At early steps of the autophagy process, Zda physically interacts with both Atg1 and Vps34, likely enabling direct activation of Vps34 by Atg1 phosphorylation (Melani et al., 2017). The final steps of exocytosis, from granule docking to granule fusion with the plasma membrane, require the concatenated action of dozens of proteins (Wang, Zhang, & Xiao, 2019). Thus, it is conceivable that a molecular platform on which these actions are sequentially organized is required, and Zda could provide such molecular platform. The Zda mammalian orthologue FKBP8 interacts with the apoptosis regulator Bcl-2, and this interaction depends on the formation of a complex with Ca^++^/CaM (Edlich et al., 2005). Therefore, since Ca^++^ is essential for GG-APM fusion (Wang et al., 2019), it seems likely that the activity of Zda in exocytosis is modulated by Ca^++^.

It was previously shown that Rho1 is recruited to GGs, presumably from the apical membrane, after the GGs fuse with the APM to induce formation of the acto-myosin coat around GGs (Rousso et al., 2016). Based on the analysis of fusion markers, we conclude that Zda is required for GG fusion with the APM, and thus, Zda probably operates upstream of Rho1. Our genetic screen identified the small GTPase RalA as the only gene, out of 64 candidates analyzed, whose knock down phenotype resembles that of Zda loss of function. RalA is a small GTPase of the Ras superfamily believed to participate in tethering exocytic vesicles to the plasma membrane through its interaction with the exocyst complex (X. W. Chen, Leto, Chiang, Wang, & Saltiel, 2007; Holly, Mavor, Zuo, & Blankenship, 2015; Jin et al., 2005; Teodoro et al., 2013). Genetic interaction experiments support the notion that Zda operates downstream of RalA. Given the previously described function of RalA in exocytosis, as a mediator of docking or anchoring of exocytic vesicles to the plasma membrane (Holly et al., 2015; Teodoro et al., 2013), our data therefore suggest that Zda probably operates downstream of the docking step, and before membrane fusion.

Overall, we have defined the immunophilin Zda as a novel player in the process of regulated exocytosis in different organs throughout development. Zda does not play a role in GG biogenesis, and we have narrowed down its window of action to the steps that precede GG-APM fusion. Even though further research is required to define its precise molecular function, we propose that Zda might operate as a molecular platform where different molecules involved in the fusion process interact with each other, and that this activity requires Ca^++^/CaM binding.

## EXPERIMENTAL PROCEDURES

### Fly stocks and genetics

All fly stocks and crosses were kept on standard corn meal/agar medium at 25 °C, except for the crosses involving RNAi that were kept at 29°C. Crosses were set up in vials containing 5 males and five females of the required genotypes. Crosses were flipped every 24 hours to avoid larval overcrowding. Embryo collecting cages were set up using 40-50 females and 10-20 males of the required genotypes. Agar plates were changed every 12 hours, and plates were left at 25C for another 12 hours, and 1^st^ instar larvae of the desired genotypes were sorted to vials to allow larval development.

The following *D. melanogaster* lines were from the Bloomington *Drosophila* Stock Center (http://flystocks.bio.indiana.edu): *sgs3-GFP* (BL5884), *sgs3-GFP* (BL5885), *UAS-white*^*RNAi*^ (BL33613), *actin-Gal4* (BL4414), *forkhead-Gal4* (BL78060), *UAS-lifeact-ruby* (BL35545), *UAS-dicer2* (BL24650), *UAS-PLCγ-PH-EGFP* (BL58362), *zda*^*trojan*^ (BL77787), *dilp2-Gal4* (BL37516). *UAS-zda^RNAi^* (v106020) was obtained from the Vienna Drosophila RNAi Center (https://stockcenter.vdrc.at). *zda*^*null*^ and UAS-mCh-Zda were previously reported (Melani et al., 2017); *sgs3-dsRed* was generated by A. Andres’ Lab (Costantino et al., 2008); *P0206-Gal4* and *phantom-Gal4* were gifts from P. Leopold. UAS-RNAi lines used in the screen (Supplementary Table 1) were obtained from the Bloomington *Drosophila* Stock Center (http://flystocks.bio.indiana.edu) or from the Vienna Drosophila Stock Center (https://stockcenter.vdrc.at).

### Cloning and transgenic line generation

cDNA encoding full length *Zda* or its truncated versions (deletion of aminoacids 190-320 = Zda^ΔCaM/ΔTPR^; deletion of aminoacids 375-395 = Zda^Δ™^) were cloned into ENTR-mCherry vector using Eco-RI site on N-terminal and Not-I on C-terminal; then gateway into the pUAST plasmids. The plasmids pUAST-mCherry-Zda-ΔCaM/ΔTM were used to generate transgenic flies. Transformants were produced by BestGene inc. (Chino Hills, CA, USA) using methodology based on procedures described previously (Rubin & Spradling, 1982).

### Tissue staining, visualization and image processing

Tissues were dissected in PBS (137 mM NaCl, 2.7 mM KCl, 4.3 mM Na2HPO4, 1.47 mM KH2PO4, [pH 8]) and either fixed in 4% methanol-free formaldehyde for 30 minutes at room temperature, or imaged directly under confocal microscope. For filamentous actin staining, fixed tissues were incubated for 2 hours with Alexa Fluor 546 Phalloidin (ThermoFisher Scientific 1:400) in PBS-0.1% Triton X-100 (PT). When needed, 300 nM 4′,6-diamidine-2-phenylindole (DAPI) was simultaneously added. For antibody staining, fixed tissues were washed three times in PT and blocked with PT bovine serum albumin 5%. Primary antibodies were incubated overnight at 4°C followed by 3 washes with PT, and 4-hour incubation with fluorophore-conjugated secondary antibodies. Primary antibodies used in this study were guinea pig anti-PTTH (1:400,– (Yamanaka et al., 2013)), rat anti-Dilp2 (1:400, Ref. (Géminard et al., 2006)), gift from Leopold’s Lab, and anti-GFP (1:10.000, chicken, Sigma G6539). Secondary antibodies used were Alexa Fluor 488 anti-chicken (1:400, Thermo Fisher Scientific), Alexa Fluor 546 anti-rat (1;400), Alexa Fluor 648 anti-guinea pig (1:400). Stained tissues were mounted in gelvatol mounting medium (Sigma) and imaged.

Images were captured using a Carl Zeiss confocal microscope LSM 710 with a Plan-Apochromat 63X/1.4NA oil objective, or a Carl Zeiss LSM 880 with a Plan-Apochromat 20X/0.8NA, or a Leica SP5 DS 40X objective. Insets of Figure 2E, were captured using Airyscan superresolution and Z-stacks were imaged in 100nm steps, pixel resolution of 1532×1532, and reconstructed using Zen-Zeiss software. Images were processed using ImageJ (NIH, Bethesda, MD) according to adjust contrast and/or merge files.

### Developmental Timing Curves

Developmental timing experiments were done at 25°C. Three to four-hour time cuts of embryos laid on apple juice plus yeast paste plates were aged for 20 hours at which point freshly ecdysing L1 larvae were transferred to vials. Each vial contained 45 L1s as indicated in each experiment. The time until pupariation was scored every 6 hours; data from at least 3 vials were compiled.

### Rescue by 20-hydroxyecdysone feeding

Four-hour egg collections were made on agar plates, and after 20 hours L1 larvae were collected and grown at a density of 40 animals per vial at 18°C. At 3^rd^ larval instar, larvae were transferred to fresh vials, and maintained 25°C until pupariation. The latter vial was supplemented or not with 20-hydroxyecdysone every 12 h (Cayman Chemical, dissolved in 95% ethanol, final concentration of 0.2 mg/mL) until puparium formation. The time of pupariation was scored as above (Developmental Timing Curves).

### Raising L2 Larvae for Timed Sample Collections for Ecdysone titers

Timed samples were raised at 25°C. Forty newly-ecdysed L1s, precisely timed on apple juice collection plates, were transferred at 1-hour intervals to 35 mm plates containing agar with surface granules of live baker’s yeast, and let them develop for ~20 hours. At this time point, larvae were monitored for morphological features of second instar to assess ecdysis. Two hour collections of freshly ecdysed L2s were transferred to fresh plates and let them develop until L2-L3 ecdysis, ~24 h for phm>dcr2 and ~36 h for phm>zda-RNAi dcr2. Staged larvae were removed from the medium, washed twice in water, dried on a Kimwipe, and stored at −80.

### Ecdysone Titers

Biological replicates of 40-60 larvae were homogenized twice in methanol and cleared by centrifugation and brought to a final volume of 450 µl. Duplicate samples were dried and resuspended in 50 µl EIA buffer (spi bio 20-HE ELISA kit – A05120), and measured with a Spi Bio Kit for 20-HE ELISA following manufacturer’s recommendations. The standard curve was built using GraphPad Prism software, non-linear regression curve fit.

### Pupal size determination

For pupal volume estimation 20-30 1^st^ instar larvae of the desired genotype were transferred to food vials with 4% corn meal and grown at 29°C, and pupae were photographed under dissection microscope. Pupal length (L) and diameter (D) were measured using ImageJ, and pupal volume was calculated as previously reported (Galagovsky et al., 2014).

### Dilp2 quantification

For Dilp2 quantification in IPCs, larvae were sorted 24h after egg lay, and 40 individuals of the desired genotype transferred to vials with 4% corn meal and grown at 29°C. Early 3^rd^ instar larvae before reaching the critical weight, were transferred to agar plates for 14-16h (starvation treatment), brains were dissected in PBS, and fixed in methanol free formaldehyde 4% for 30 minutes at room temperature,then washed 3 times for 15 min with 0,3% Triton X-100 in PBS, and blocked in 5% BSA, 0,3% Triton X-100 in PBS for 2hs. Samples were then incubated with rat anti-dilp2 (1:500) (kind gift of Pierre Leopold) overnight at 4°C, and then with an Alexa 647 anti-rat secondary antibody (Sigma 1:250). Stained samples were mounted in gelvatol mounting medium (Sigma) and imaged under a Carl Zeiss LSM 880 confocal microscope with a Plan-Apochromat 20X/0.8NA objective, with 8-bit color depth, 2,5x digital zoom and pixel resolution of 1024 × 1024. Z-stacks were imaged in 2,97μm steps over a total depth of 32,16 μm. Fluorescence quantification was assessed using ImageJ. Fluorescence intensities from all the slices were summed, and areas were selected based on the channel showing IPCs. Maximum Z-projections were made with Zen-Zeiss software.

### Sgs3-GFP retention phenotype

Larvae or prepupae of the desired genotype and developmental stage were visualized and photographed inside vials under a fluorescence dissection microscope. Each experiment was repeated at least 3 times.

### Statistical analyses

Statistical significance was calculated using the two-tailed Student’s t test when comparing two values, and one-way analysis of variance (ANOVA), followed by a Tuckey’s test with a 95% confidence interval (p < 0.05) when comparing multiple values. When needed, Grubb’s test was used to identify the values that were significant outliers from the rest (p < 0.05) (https://graphpad.com/quickcalcs/grubbs2/). In all cases, error bars represent the SD.

## ACKNOWLEDGMENTS

We are grateful to Dr. Andrew Andres, Dr. Gabor Juhasz, the Bloomington Stock Centre and the Vienna Drosophila Resource Centre for fly strains. We thank Dr. Pierre Leopold for sharing the anti-Dilp2 antibody. We thank all members of the Wappner lab for discussions, Dr. Andrés Rossi for technical support with confocal microscopy, Andrés Liceri for fly food preparation and FIL personnel for assistance.

---

**Supplementary Figure 1.**
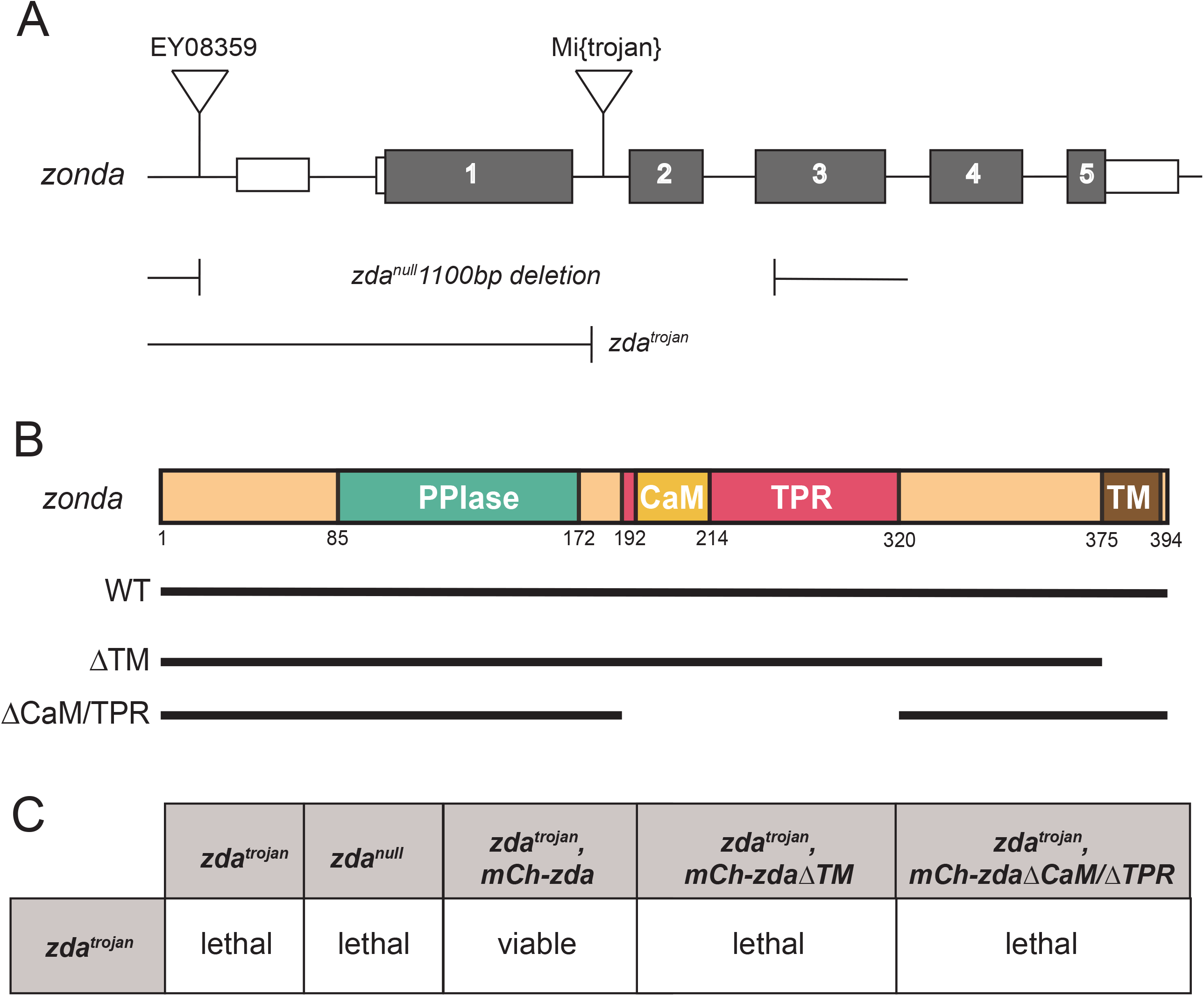
(A)The *zda* locus (*CG5482*) encompasses 5 exons. Imprecise excision of a P element (*EY08359*) inserted in the 5’UTR of the gene was induced, and generated a 1100 base pairs deletion giving rise to the *zda*^*null*^ allele. The *Mi{Trojan-Gal4.0}zda[MI07788-TG4.0]* insertion at the 2^nd^ intron generates a truncated version of Zda. (B) Schematic representation of Zda predicted domains. From the N- to the C-terminus: Peptidyl prolyl cis/trans isomerase (PPIase), Calmodulin binding (CaM), Tetratricopeptide repeat (TPR), and Transmembrane (TM) domains are depicted. Deletion of specific domains used to generate transgenic lines are shown below. (C) Complementation test. Analysis of viability to adulthood of different combinations *zda* alleles.

**Supplementary Figure 2.**
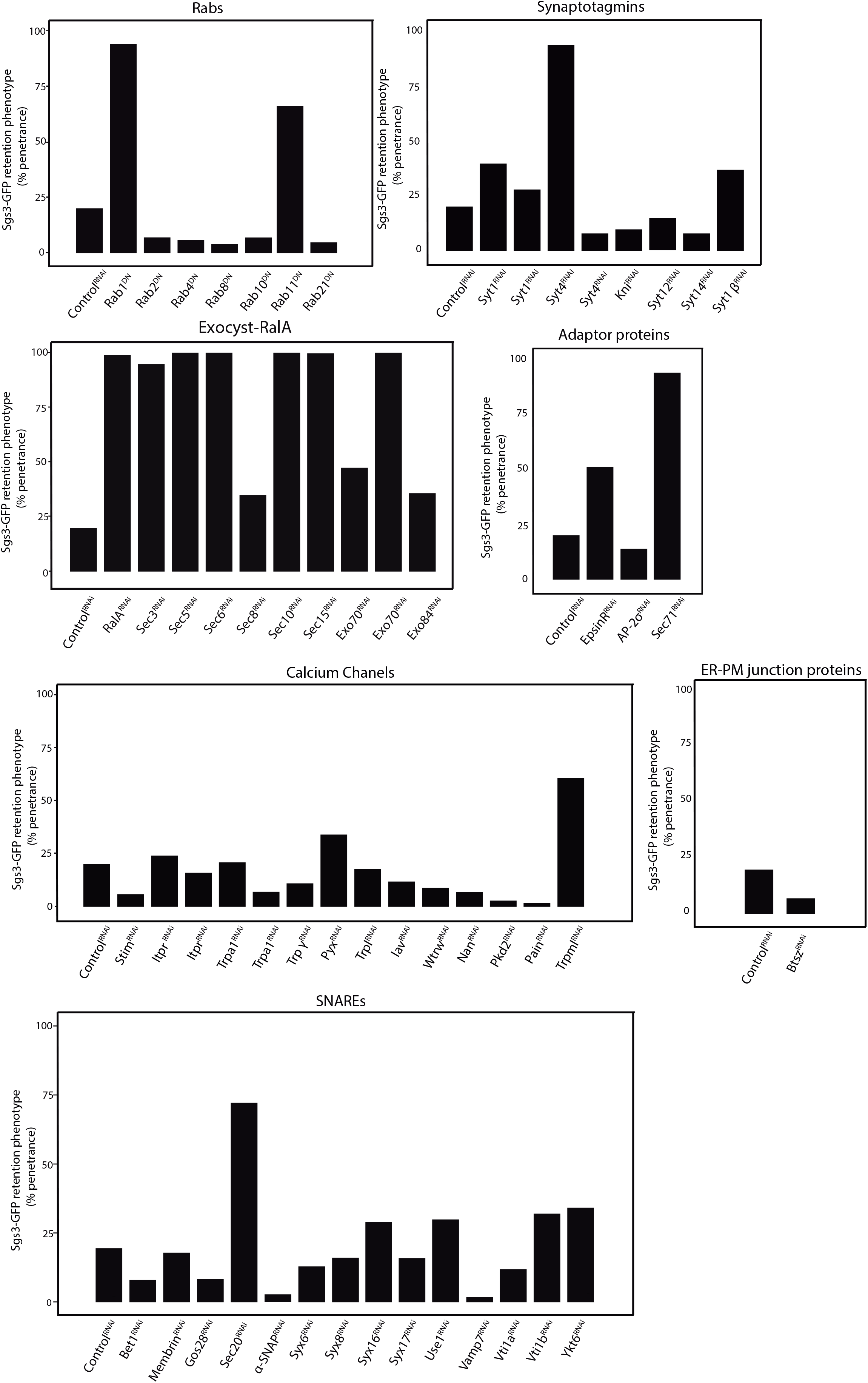
Quantification of Sgs3-GFP retention inside salivary glands of prepupae after loss of function of the indicated genes. RNAi or dominant negative constructs where expressed in salivary glands using a *fkh-Gal4* driver. The penetrance of the phenotypes is depicted. The number of individuals scored in each case is shown in Supplementary Table 1.

**Supplementary Figure 3.**
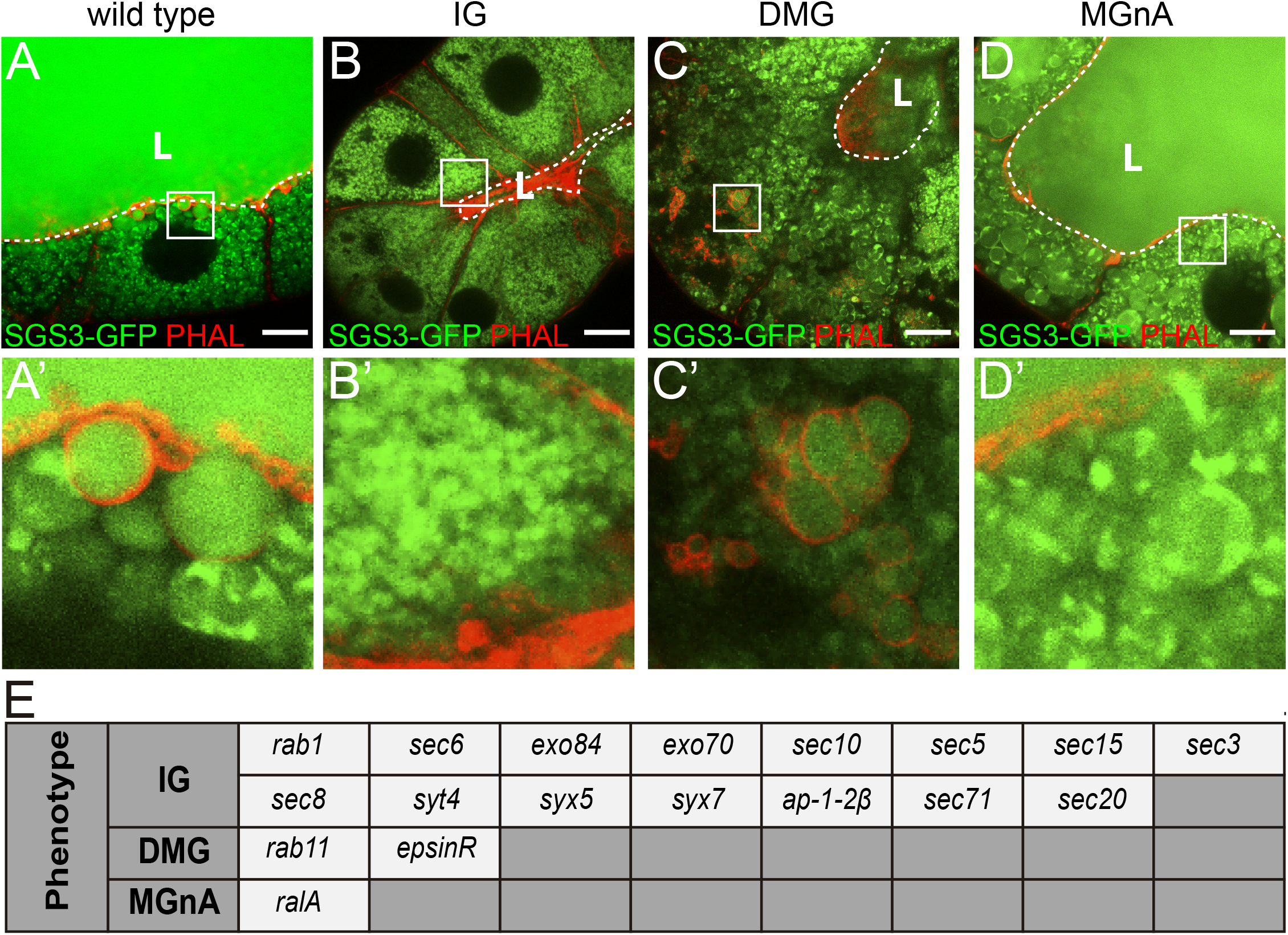
Phenotypic categories identified at the secondary screen: A) wild type; B) Small GGs; C) delocalized GGs (DGG); D) Mature GGs without actin mesh (MGGnA). GGs are labeled with Sgs3-GFP (green), and filamentous actin is labelled with phalloidin (red); “L” indicates the lumen that is also marked with a dashed line. Scale bar: 20μm. Crop size 20μm×20μm. (E) The genes identified in each category are listed.

**Supplementary Table 1.**
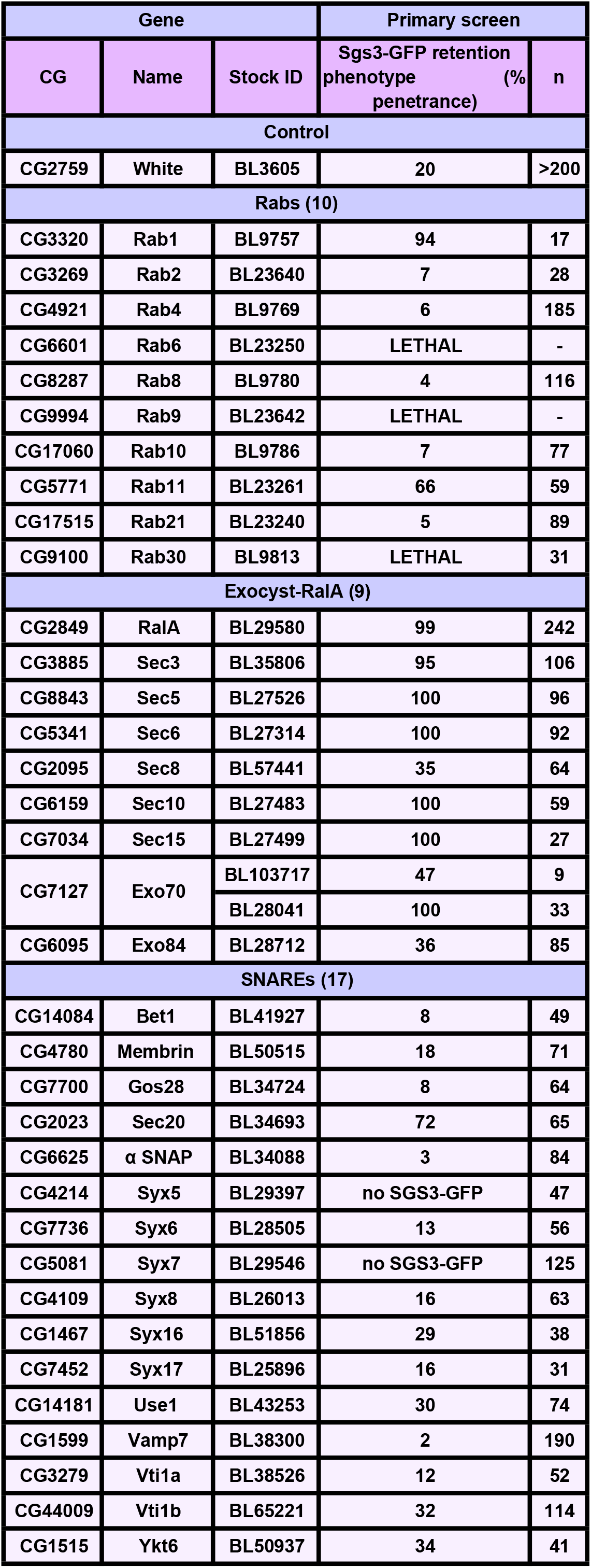
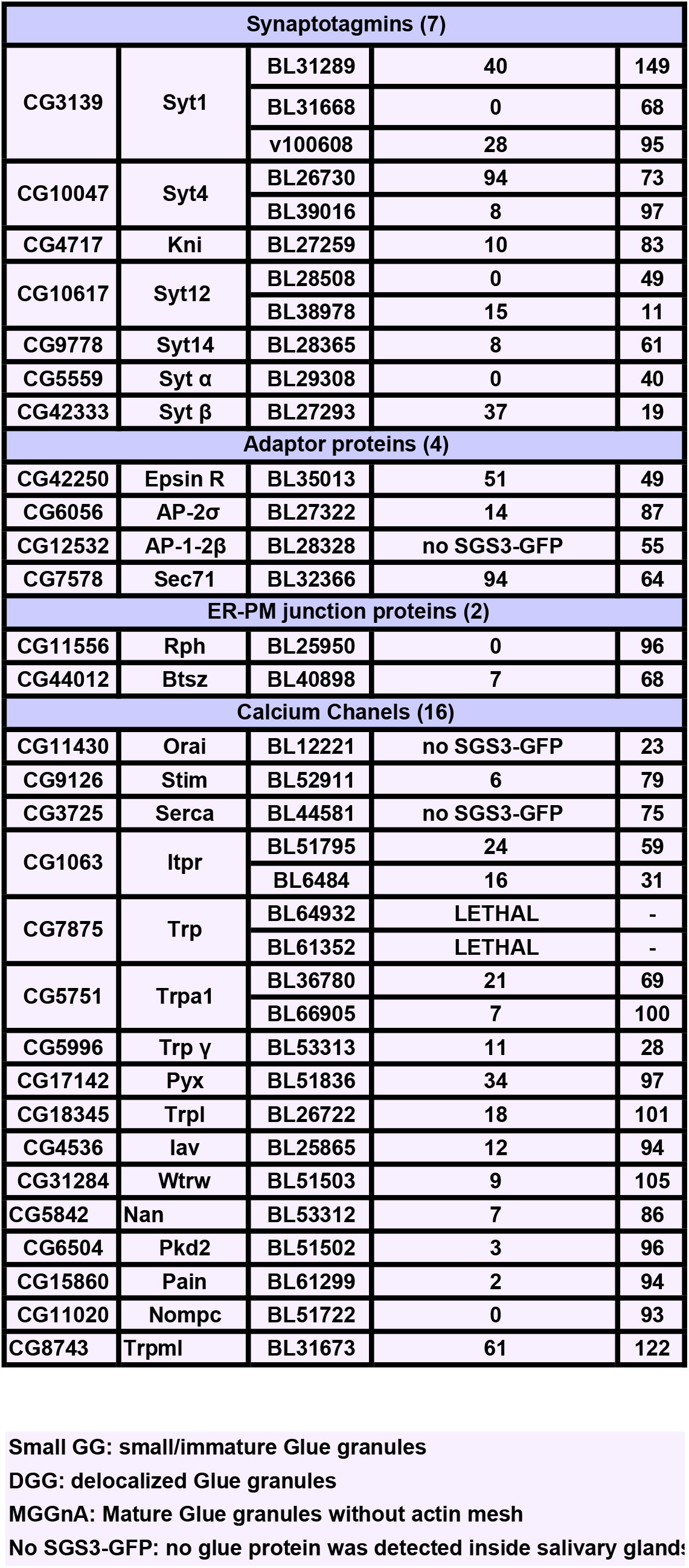
List of genes screened for Sgs3-GFP exocytosis.

| Gene |  |  | Primary screen |  |
| --- | --- | --- | --- | --- |
| CG | Name | Stock ID | Sgs3-GFP retention phenotype (% penetrance) | n |
| Control |  |  |  |  |
| CG2759 | White | BL3605 | 20 | >200 |
| Rabs (10) |  |  |  |  |
| CG3320 | Rab1 | BL9757 | 94 | 17 |
| CG3269 | Rab2 | BL23640 | 7 | 28 |
| CG4921 | Rab4 | BL9769 | 6 | 185 |
| CG6601 | Rab6 | BL23250 | LETHAL | - |
| CG8287 | Rab8 | BL9780 | 4 | 116 |
| CG9994 | Rab9 | BL23642 | LETHAL | - |
| CG17060 | Rab10 | BL9786 | 7 | 77 |
| CG5771 | Rab11 | BL23261 | 66 | 59 |
| CG17515 | Rab21 | BL23240 | 5 | 89 |
| CG9100 | Rab30 | BL9813 | LETHAL | 31 |
| Exocyst-RalA (9) |  |  |  |  |
| CG2849 | RalA | BL29580 | 99 | 242 |
| CG3885 | Sec3 | BL35806 | 95 | 106 |
| CG8843 | Sec5 | BL27526 | 100 | 96 |
| CG5341 | Sec6 | BL27314 | 100 | 92 |
| CG2095 | Sec8 | BL57441 | 35 | 64 |
| CG6159 | Sec10 | BL27483 | 100 | 59 |
| CG7034 | Sec15 | BL27499 | 100 | 27 |
| CG7127 | Exo70 | BL103717 | 47 | 9 |
|  |  | BL28041 | 100 | 33 |
| CG6095 | Exo84 | BL28712 | 36 | 85 |
| SNAREs (17) |  |  |  |  |
| CG14084 | Bet1 | BL41927 | 8 | 49 |
| CG4780 | Membrin | BL50515 | 18 | 71 |
| CG7700 | Gos28 | BL34724 | 8 | 64 |
| CG2023 | Sec20 | BL34693 | 72 | 65 |
| CG6625 | $\alpha$ SNAP | BL34088 | 3 | 84 |
| CG4214 | Syx5 | BL29397 | no SGS3-GFP | 47 |
| CG7736 | Syx6 | BL28505 | 13 | 56 |
| CG5081 | Syx7 | BL29546 | no SGS3-GFP | 125 |
| CG4109 | Syx8 | BL26013 | 16 | 63 |
| CG1467 | Syx16 | BL51856 | 29 | 38 |
| CG7452 | Syx17 | BL25896 | 16 | 31 |
| CG14181 | Use1 | BL43253 | 30 | 74 |
| CG1599 | Vamp7 | BL38300 | 2 | 190 |
| CG3279 | Vti1a | BL38526 | 12 | 52 |
| CG44009 | Vti1b | BL65221 | 32 | 114 |
| CG1515 | Ykt6 | BL50937 | 34 | 41 |

| Synaptotagmins (7) |  |  |  |  |
| --- | --- | --- | --- | --- |
| CG3139 | Syt1 | BL31289 | 40 | 149 |
|  |  | BL31668 | 0 | 68 |
|  |  | v100608 | 28 | 95 |
| CG10047 | Syt4 | BL26730 | 94 | 73 |
|  |  | BL39016 | 8 | 97 |
| CG4717 | Kni | BL27259 | 10 | 83 |
| CG10617 | Syt12 | BL28508 | 0 | 49 |
|  |  | BL38978 | 15 | 11 |
| CG9778 | Syt14 | BL28365 | 8 | 61 |
| CG5559 | Syt $\alpha$ | BL29308 | 0 | 40 |
| CG42333 | Syt $\beta$ | BL27293 | 37 | 19 |
| Adaptor proteins (4) |  |  |  |  |
| CG42250 | Epsin R | BL35013 | 51 | 49 |
| CG6056 | AP-2σ | BL27322 | 14 | 87 |
| CG12532 | AP-1-2β | BL28328 | no SGS3-GFP | 55 |
| CG7578 | Sec71 | BL32366 | 94 | 64 |
| ER-PM junction proteins (2) |  |  |  |  |
| CG11556 | Rph | BL25950 | 0 | 96 |
| CG44012 | Btsz | BL40898 | 7 | 68 |
| Calcium Channels (16) |  |  |  |  |
| CG11430 | Orai | BL12221 | no SGS3-GFP | 23 |
| CG9126 | Stim | BL52911 | 6 | 79 |
| CG3725 | Serca | BL44581 | no SGS3-GFP | 75 |
| CG1063 | Itpr | BL51795 | 24 | 59 |
|  |  | BL6484 | 16 | 31 |
| CG7875 | Trp | BL64932 | LETHAL | - |
|  |  | BL61352 | LETHAL | - |
| CG5751 | Trpa1 | BL36780 | 21 | 69 |
|  |  | BL66905 | 7 | 100 |
| CG5996 | Trp $\gamma$ | BL53313 | 11 | 28 |
| CG17142 | Pyx | BL51836 | 34 | 97 |
| CG18345 | Trpl | BL26722 | 18 | 101 |
| CG4536 | Iav | BL25865 | 12 | 94 |
| CG31284 | Wtrw | BL51503 | 9 | 105 |
| CG5842 | Nan | BL53312 | 7 | 86 |
| CG6504 | Pkd2 | BL51502 | 3 | 96 |
| CG15860 | Pain | BL61299 | 2 | 94 |
| CG11020 | Nompc | BL51722 | 0 | 93 |
| CG8743 | Trpml | BL31673 | 61 | 122 |

Small GG: small/immature Glue granules
DGG: delocalized Glue granules
MGGnA: Mature Glue granules without actin mesh
No SGS3-GFP: no glue protein was detected inside salivary glands

